# Reliable longitudinal brain age prediction in stroke patients: Associations with cognitive function and response to cognitive training

**DOI:** 10.1101/687079

**Authors:** Geneviève Richard, Knut Kolskår, Kristine M. Ulrichsen, Tobias Kaufmann, Dag Alnæs, Anne-Marthe Sanders, Erlend S. Dørum, Jennifer Monereo Sánchez, Anders Petersen, Hege Ihle-Hansen, Jan Egil Nordvik, Lars T. Westlye

## Abstract

Cognitive deficits are important predictors for outcome, independence and quality of life after stroke, but often remain unnoticed and unattended because other impairments are more evident. Computerized cognitive training (CCT) is among the candidate interventions that may alleviate cognitive difficulties, but the evidence supporting its feasibility and effectiveness is scarce, partly due to the lack of tools for outcome prediction and monitoring. Magnetic resonance imaging (MRI) provides candidate markers for disease monitoring and outcome prediction. By integrating information not only about lesion extent and localization, but also regarding the integrity of the unaffected parts of the brain, advanced MRI provides relevant information for developing better prediction models in order to tailor cognitive intervention for patients, especially in a chronic phase.

Using brain age prediction based on MRI based brain morphometry and machine learning, we tested the hypotheses that stroke patients with a younger-appearing brain relative to their chronological age perform better on cognitive tests and benefit more from cognitive training compared to patients with an older-appearing brain. In this randomized double-blind study, 54 patients who suffered mild stroke (>6 months since hospital admission, NIHSS<7 at hospital discharge) underwent 3-weeks CCT and MRI before and after the intervention. In addition, patients were randomized to one of two groups receiving either active or sham transcranial direct current stimulation (tDCS). We tested for main effects of brain age gap (estimated age – chronological age) on cognitive performance, and associations between brain age gap and task improvement. Finally, we tested if longitudinal changes in brain age gap during the intervention were sensitive to treatment response. Briefly, our results suggest that longitudinal brain age prediction based on automated brain morphometry is feasible and reliable in stroke patients. However, no significant association between brain age and both performance and response to cognitive training were found.

## Introduction

Stroke is among the most common causes of acquired cognitive disabilities during adulthood, with a projected increase in prevalence over the next decades due to the aging population (Feigin et al. 2014; Feigin et al. 2017). Despite recent reductions in stroke-related mortalities, largely due to major improvements in acute health care and treatment (Zhang et al. 2012) many stroke survivors suffer from long-term and pervasive cognitive deficits (Barbay et al. 2018; Barker-Collo et al. 2010; Cumming et al. 2014; Haacke et al. 2006; Nakling et al. 2017; Patel et al. 2002) that often remain unnoticed by the health care system due to its typically delayed manifestation (Jacova et al. 2012; Kalaria et al. 2016).

Previous studies and treatment programs have largely targeted patients in the acute and sub-acute phase, as it has been assumed that recovery and cognitive rehabilitation are more likely to be successful during a limited time window following the insult (Zucchella et al. 2014). Whereas the temporal aspects of cognitive interventions following stroke is important, evidence suggests that recovery can also occur in chronic stages, i.e. years after the insult (Berthier et al. 2011; Moss & Nicholas 2006). As a result, there is an increasing need for developing and validating tools that can be used to predict long-term outcome and for monitoring of the effects of cognitive rehabilitation after stroke (Hope et al. 2013).

Advanced neuroimaging techniques based on magnetic resonance imaging (MRI) offer a range of candidate markers for disease monitoring and outcome prediction. In addition to providing detailed information about the localization and extent of the lesion, which represent key clinical information in the acute phase, imaging techniques allow for a characterization of the structural and functional integrity of the whole brain, including areas not directly damaged by the stroke (Kalaria et al. 2016; Werden et al. 2017). This information is highly relevant in a cognitive rehabilitation context, where the potential for improvement and recovery are not only defined by the lesion itself, but by the integrity and efficiency of the unaffected brain regions (Ihle-Hansen et al. 2014). Further, it is widely acknowledged that the brain systems supporting cognitive functions are broadly distributed, supporting a network-based conceptualization of the functional neuroanatomy of cognitive functions. Hence, lesions in widely different parts of the brain may result in overlapping cognitive symptoms, depending on the brain networks involved (Guggisberg et al. 2019). A direct implication of this is that both the degree of cognitive impairment and the individual potential for improvement in response to intervention may be less dependent on the exact characteristics of the lesion than the structural integrity of the unaffected brain networks.

Here, we test this concept by utilizing multivariate brain age prediction using machine learning and sensitive measures of brain morphometry. Briefly, combining a wide array of informative brain imaging features in a prediction model allows for an accurate prediction of the age of an unseen individual (Franke et al. 2012; Franke et al. 2010). The degree to which the model under- or over-estimate the individual’s age has been shown to be sensitive to a variety of health-related characteristics, including cognitive function and mortality (Boyle et al. 2019; Cole & Franke 2017; Cole et al. 2018; Richard et al. 2018), and brain age prediction using MRI data has recently been shown to be sensitive both to the clinical manifestation and polygenic risk of various brain disorders (Høgestøl et al. 2019a; Kaufmann et al. 2018).

Based on the notion that brain age prediction offers a sensitive summary measure of brain health and integrity, we first tested whether brain age is sensitive to cognitive function in chronic stroke patients. Next, to assess the predictive value of brain age prediction in a cognitive rehabilitation context, we tested if brain age prior to the intervention is associated with response to an intensive computerized cognitive training (CCT) program. As a follow-up analysis to a previous study (Kolskår et al. 2019) reporting no robust beneficial effects of transcranial direct brain stimulation (tDCS) on cognitive improvement, we assessed if any beneficial effects of tDCS (active vs sham) would be dependent on brain age. Finally, we tested to which degree longitudinal changes in brain age during the course of the intervention are sensitive to treatment response. We hypothesized that (1) brain age prediction would constitute a reliable and sensitive method for characterizing individual level brain health. We further anticipated that (2) patients with a relatively low brain age (which may imply higher cognitive or brain reserve) would show better cognitive function at baseline, and (3) would show larger improvements in task performance. Lastly, to the extent that intensive cognitive training shows beneficial effects on cognitive performance and the brain (Engvig et al. 2010), we hypothesized that (4) cognitive gains would be reflected in longitudinal changes in brain age during the course of the intervention.

We tested these hypotheses in a group of 54 chronic patients who suffered mild stroke (> 6 months since hospital admission, NIHSS < 7 at hospital discharge) invited to take part in a randomized, double blind study aimed to test the utility of tDCS in combination with CCT to improve cognitive performance following stroke (Kolskår et al. 2019; Ulrichsen et al. 2019). For unbiased brain age prediction, we utilized a large independent training set, and employed stringent procedures for multiple comparison correction to increase the robustness of the results.

## Materials and methods

Table 1 summarizes key clinical and demographic information for the patient group. Patients were recruited with the main aim of testing the clinical feasibility of combining CCT and tDCS to improve cognitive function in chronic stroke patients. Description of the extent and localization of individual patient lesions, as well as recruitment procedures are detailed in (Kolskår et al. 2019). Briefly, patients admitted to the Stroke Unit at Oslo University Hospital and at Diakonhjemmet Hospital, Oslo, Norway during 2013-2016 were invited to participate through letters. Stroke was defined as any form of strokes of either ischemic or hemorrhagic etiology; transient ischemic attacks (TIA) were excluded. Additional exclusion criteria included MRI contraindications and other neurological diseases diagnosed prior to the stroke.

**Table 1.**
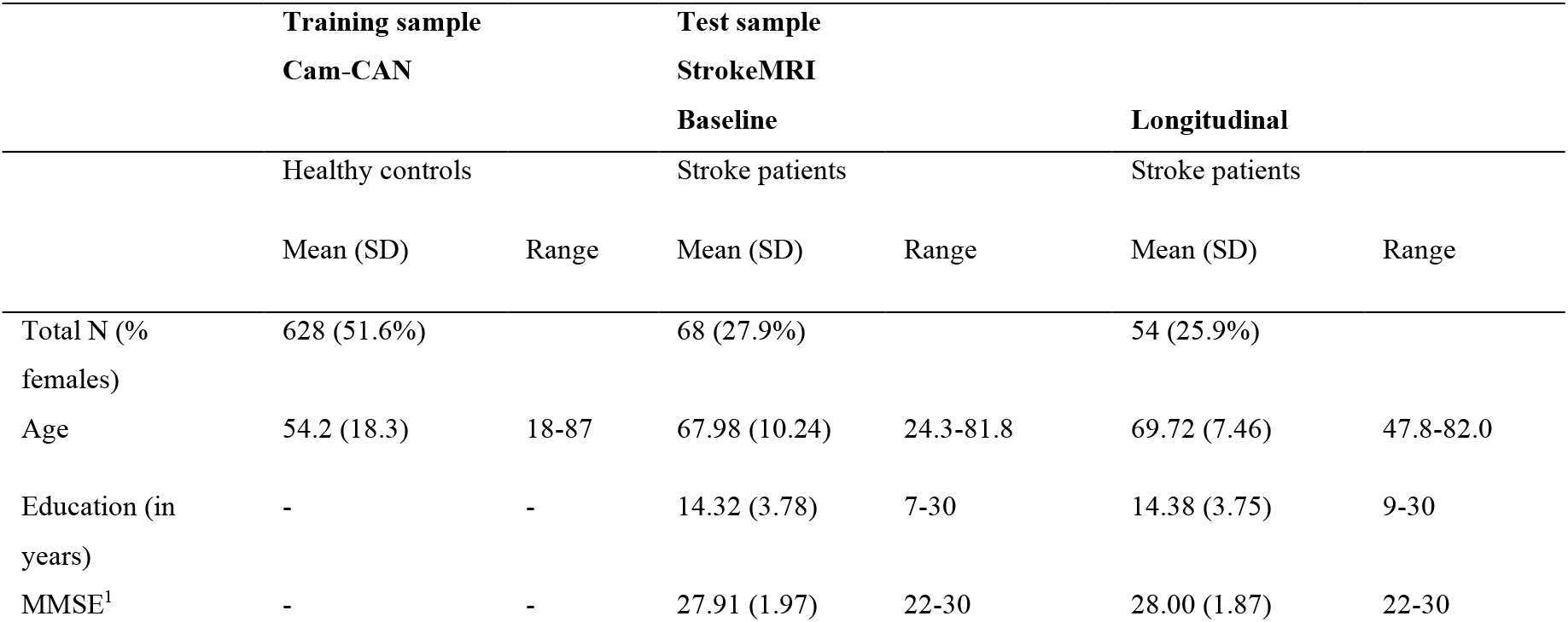
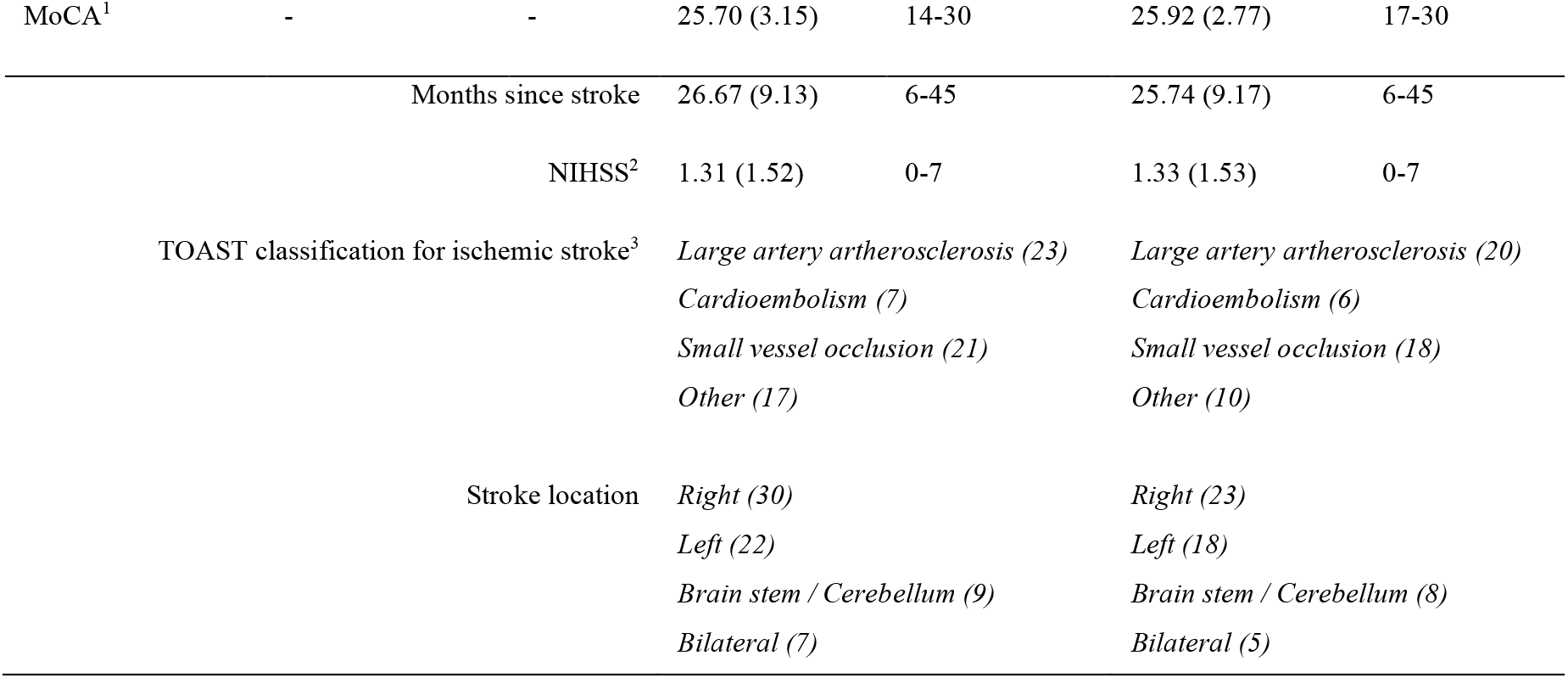
Demographics and sample characteristics. ^1^MMSE and MoCA scores at inclusion. ^2^NIHSS score at hospital discharge. ^3^One patient had intracerebral hemorrhage (Kolskår et al. 2019; Ulrichsen et al. 2019).

Approximately 250 patients responded to the letter, of which 72 completed the first assessment and 54 patients completed the full protocol; including three MRI brain scan sessions, three sessions with cognitive assessments, one EEG assessment, and seven CCT sessions in addition to 10 CCT sessions performed at home.

Four patients were excluded from the analysis in the current study. Two were excluded based on poor quality or incomplete MRI data, one based on incomplete cognitive assessment at baseline and one due to lack of confirmed stroke. The remaining 68 patients were included in the brain age estimation and associations with baseline cognitive performance (age = 24.3-81.8, mean = 67.98, SD = 10.24, 19 females). All 54 patients who completed the training sessions were included in the remaining analyses (age = 47.8-82.0, mean = 69.72, SD = 7.46, 14 females).

### Training set for brain age prediction

The healthy controls used as training set for the age prediction model were obtained from the Cambridge Centre for Ageing and Neuroscience (Cam-CAN) sample (http://www.mrc-cbu.cam.ac.uk/datasets/camcan/; (Shafto et al. 2014; Taylor et al. 2017)). Briefly, volunteers were recruited to Cam-CAN through a large-scale collaborative research project funded by the Biotechnology and Biological Sciences Research Council (BBSRC, grant number BB/H008217/1), the UK Medical Research Council and University of Cambridge. For more information, see http://www.cam-can.org. Data from 628 individuals (age = 18-87, mean = 54.2, SD = 18.3, 324 females) were included in the training set (Richard et al. 2018).

### Cognitive assessment at baseline

Similar to our recent study (Richard et al. 2018), cognitive performance at baseline was assessed with a set of neuropsychological and computerized tests assumed to be sensitive to cognitive aging, including the Montreal Cognitive Assessment (MoCA; Nasreddine et al. 2005), the vocabulary and matrix subtests of the Wechsler Abbreviated Scale of Intelligence (WASI; Wechsler 1999), the California Verbal Learning Test (CVLT-II; Delis et al. 2000), and the Delis-Kaplan Executive Function System (D-KEFS) color word interference test (Stroop; Delis et al. 2001). We included several computerized tests from the Cognitive Assessment at Bedside for iPAD (CABPad; Willer et al. 2016), including motor speed, verbal fluency (phonological and semantic), working memory (forward and backward memory span), spatial Stroop (executive control of attention), spatial attention span, and symbol digit coding tests. Further, a computerized test based on the Theory of Visual Attention (TVA; Bundesen 1990; Bundesen & Habekost 2008; Dyrholm et al. 2011) provided measures of visual short-term memory capacity (*K*), processing speed (*C*), and perceptual threshold (*t*_0_). Several variables were highly correlated, and we used the clustering solution from Richard et al. (2018), which included seven broad cognitive domains. Cluster 1 reflected memory and learning (CVLT, attention span, MoCA), cluster 2 visual processing speed (TVA-parameters *C* and *t*_0_), cluster 3 verbal skills (phonological and semantic flow), cluster 4 attentional control and speed (spatial Stroop), cluster 5 executive control and speed (color-word Stroop), cluster 6 reasoning and psychomotor speed (matrix, symbol coding and motor speed, visual short-term memory capacity (TVA-parameter *K*)), and cluster 7 working memory (forward and backward memory span). Briefly, the clusters were computed using normalized sum scores of highly correlated test scores. Prior to calculating summary scores based on the seven clusters mentioned above, we used *outlierTest* from the car package (Fox & Weisberg 2011) to identify the most extreme observations based on a linear model, including age and sex. 17 observations were identified as outliers based on a Bonferroni corrected p < 0.05 and treated as missing values, we then replaced these extremes and imputed the 75 missing values (2.63% of the scores were missing/incomplete) using predictive mean matching *(pmm)* method from the *mice* package in R (multivariate imputation by chained equations; Buuren & Groothuis-Oudshoorn 2011).

### CCT protocol

All patients completed a computerized working memory training program consisting of 25 online training sessions (Cogmed Systems AB, Stockholm, Sweden). Similar to our recent study (Kolskår et al. 2019), we used data from 17 of the 25 training sessions over a period of three to four weeks, corresponding to approximately five weekly training sessions. Seven sessions were carried out at the hospital, of which six were in combination with tDCS (either sham or active stimulation). On average, patients received two training sessions with tDCS per week with a minimum of one day between each session. The remaining 10 training sessions were home-training. Each training session took approximately 45 minutes in which the participant completed eight different exercises. In total, 10 different tasks targeting verbal and visuospatial working memory were used, i.e. Grid, Hidden, Cube, Sort, Digits, 3D Cube, Twist, Assembly, Rotating and Chaos. The difficulty level of each task is adapted to the participant’s performance, and in general, for each task, it takes approximately two sessions for the difficulty level to be appropriately adjusted to the individual level of performance. Thus, we discarded the two first training sessions of each task from our analysis. In addition, we included only tasks with a minimum of three training sessions after exclusion of the two first sessions, discarding Assembly and Chaos from further analysis.

### tDCS protocol

The tDCS protocol has been described in details in a prior publication (Kolskår et al. 2019). Participants were randomly assigned to an active or a sham condition, using an in-house Matlab script to randomly generate a code for each participant while ensuring that each block of 20 participants was balanced across conditions. Both the participant and the experimenter remained blinded throughout the experiment. Stimulation was delivered using a battery-driven direct current stimulator (Neuroconn DC-STIMULATOR PLUS, neuroConn GmbH, Illmenau, Germany), through 5 x 7 cm rubber pads using the following parameters: DC current = 1 mA, total duration = 20 minutes, ramp-up = 120 seconds, fade-out = 30 seconds, and current density = 28.57 μA/cm^2^. The sham stimulation followed the factory settings which include a ramp-up and a fade-out period. We used the 10-20 system for the electrode location, with the anodal electrode covering F3 and the cathodal electrode placed over O2, and fixated with rubber bands. The pads were covered with high-conductive gel (Abralyt HiCl, Falk Minow Services Herrsching, Germany) to keep the impedance threshold under < 20 kΩ. For security reason, the device has an absolute impedance threshold of 40 kΩ. Following each stimulation period, participants were asked to fill in a side-effect form. In addition, after the last stimulation session, they were asked to make a guess whether they thought they received active stimulation or sham stimulation and the reason for their guess.

### MRI acquisition

Patients were scanned on a 3T GE 750 Discovery MRI scanner with a 32-channel head coil at Oslo University Hospital. Paddings were used to reduce head motion. T1-weighted data was acquired using a 3D IR-prepared FSPGR (BRAVO) with the following parameters: repetition time (TR): 8.16 ms, echo time (TE): 3.18 ms, inversion time (TI): 450 ms, flip angle (FA): 12°, voxel size: 1 × 1 × 1 mm, field of view (FOV): 256 x 256 mm, 188 sagittal slices, scan time: 4:43 minutes.

Cam-CAN participants were scanned on a 3T Siemens TIM Trio scanner with a 32-channel head-coil at Medical Research Council (UK) Cognition and Brain Sciences Unit (MRC-CBSU) in Cambridge, UK. High-resolution 3D T1-weighted data was acquired using a magnetization prepared rapid gradient echo (MPRAGE) sequence with the following parameters: TR: 2250 ms, TE: 2.99 ms, TI: 900 ms, FA: 9°, FOV of 256 x 240 x 192 mm; voxel size =1 mm^3^ isotropic, GRAPPA acceleration factor of 2, scan time 4:32 minutes (Shafto et al. 2014).

### MRI processing

All T1-weighted images were processed using FreeSurfer 5.3 (http://surfer.nmr.mgh.harvard.edu; (Dale et al. 1999)) including brain extraction, intensity normalization, automated tissue segmentation, generation of white and pial surfaces (Dale et al. 1999). All reconstructions were visually assessed and corrected as appropriate, and data with excessive motion or other major artefacts were discarded.

For StrokeMRI, images were processed with the longitudinal Freesurfer pipeline (Reuter & Fischl 2011; Reuter et al. 2012), which substantially increases reliability and power (Reuter et al. 2012). For each individual dataset, we extracted mean cortical thickness, area and volumes from 180 regions of interests (ROIs) per hemisphere based on a surface-based atlas (Glasser et al. 2016), yielding 1080 structural brain features per individual.

### Age prediction

Based on a recent implementation (Kaufmann et al. 2018), brain age estimation was performed both using global and regional features as input. The regional brain age estimations were based on *lobesStrict* segmentation (occipital, frontal, temporal, parietal, cingulate and insulate) from Freesurfer (Dale et al. 1999). Overall, one global and 12 hemisphere specific lobe-based models were trained to estimate age in 628 healthy controls from the Cam-CAN cohort, using the same pipeline as previously described (Richard et al. 2018). We used *xgboost* package in R (extreme gradient boosting) (Chen & Guestrin 2016; Chen et al. 2017) with the following parameters: learning rate (eta) = 0.1, nround = 1500, gamma = 1, max_depth = 6, subsample=0.5, to build the prediction models. For each model, the performance was estimated using a 10-fold cross-validation procedure within the training set.

Next, we tested the performance of our trained models by predicting age in unseen subjects in the test sample. More specifically, we calculated the Pearson correlation between the predicted and the chronological, as well as the mean absolute error (MAE) in years. For each individual and for each model, we calculated the brain age gap (BAG), i.e. the difference between the estimated and chronological age, yielding 13 BAGs per individuals. Next, in order to account for age-related bias in the age prediction (Le et al. 2018), we used linear modeling to regress out the main effect of age, age^2^ and sex from each BAG, resulting in 13 residualized BAG (BAGR) used in the calculation of MAE and further analyses.

In some instances, the stroke lesions interfered with the cortical reconstruction process in Freesurfer, which inevitably influences the estimated morphometric parameters in the relevant part of the brain. In order to assess the influence of the stroke lesion on the brain age estimates, we used *outlierTest* from the *car* package (Fox & Weisberg, 2011) to identify the most extreme morphometric estimations based on a linear model, including age, age^2^ and sex. We identified 479 observations (0.24% of all observations) as extreme and replaced them using predictive mean matching (*pmm*) method from the *mice* package in R (multivariate imputation by chained equations; Buuren & Groothuis-Oudshoorn 2011). Next, we estimated brain age using the resulting data frame containing imputed estimations and compared it with the original estimations. Subsequent analyses were performed both with and without the outliers included. Briefly, the estimated brain age based on the original Freesurfer estimations and the estimations after imputing realistic values to replace outliers resulted in nearly identical outcomes. (See supplemental results for the analyses performed after removing the outliers.)

In addition to the global model including all T1 features, we calculated a robust brain age based on the median of the 12 regional brain ages.

### Processing of Cogmed data

For each participant and for each included tasks, we used linear modeling to quantify the changes in performance across the training period, i.e. the cognitive improvement, using performance as dependent variable and session number as independent variable (Kolskår et al. 2019). In addition, we used the generic function *predict* in R (Chambers & Hastie 1992) to estimate the baseline score and the final score using the resulting individual linear models for each trained task. To derive a common score across the trained tasks, we performed a principal component analysis (PCA) on the performance improvement scores and we used the first component as the individual’s performance improvement (Kolskår et al. 2019). All test scores were zero-centered and standardized prior to running the PCA.

### Statistical analysis

Statistical analyses were performed using R version 3.3.3 (2017-03-06) (R Core Team 2017). We assessed the reliability of the age estimations using intra-class coefficient (ICC) using *ICCest* function from the ICC R package (Wolak et al. 2012) across the two baseline MRI and across all three MRI sessions.

To test if patients with relative low brain age show better cognitive performance at baseline, we employed linear models with the seven summary scores based on the clustering solution from (Richard et al. 2018) as independent variable and each BAGR as the dependent variable, including age and sex as covariates. To test if patients with relative low brain age would show larger improvement in task performance, we employed linear models with Cogmed performance gain score derived from the PCA as independent variable and each BAGR as dependent variable, including age and sex as covariates. For transparency, we report both uncorrected p-values and p-values adjusted using false discovery rate (FDR; Benjamini & Hochberg 1995) from the *p.adjust* function from the stats R package (R Core Team 2017). We have previously reported no significant beneficial effects of tDCS on cognitive improvement in response to the intervention (Kolskår et al. 2019). Here, as a follow-up analysis, we added tDCS group (sham vs experimental) as an additional variable and tested for interactions between tDCS and BAGR on training gain to assess if any beneficial effects of tDCS would be dependent on BAGR.

Lastly, in order to assess if cognitive improvements in response to intensive cognitive training is associated with reduced brain age during the course of the intervention, we tested for associations between cognitive performance and BAGR by time interaction in a longitudinal context using linear mixed effects models (LME). For each trained task, we used the estimated baseline and final scores from the individual linear models, and we used BAGR from scan number 2 and 3 as timepoint one and two, respectively. Estimated task performance was entered as dependent variable, with BAGR, time, BAGR by time interaction, age and sex as fixed factors, and participant as random factor.

## Results

### Brain age predictions

Ten-fold cross-validation on the training sample (Cam-CAN) revealed relatively high correlations between chronological and predicted age for each of the 13 models, confirming reasonable model performance. Supplementary Fig. 1 shows Pearson correlation with confidence intervals between estimated brain age and chronological age within the training sample for each of the 13 trained models ranging from *r* = .84 (CI=.81-.86) for the full model to *r* = .61 (CI=.56-.66) for the model based on right cingulate features.

Table 2 shows Pearson correlation between estimated brain age and chronological age with their 95 % confidence intervals on the test sample at baseline (stroke patients) for each model, in addition to the MAE calculated from BAGR. The correlations ranged from *r* = .58 (CI=.40-.72, MAE=4.27) for the most comprehensive model based on the median of the 12 regional models to *r* = .09 (CI=-.16-.32 MAE=7.29) for the left cingulate model. See Suppl. Table 1 for the model performance after replacing outliers by imputed values.

**Table 2.**
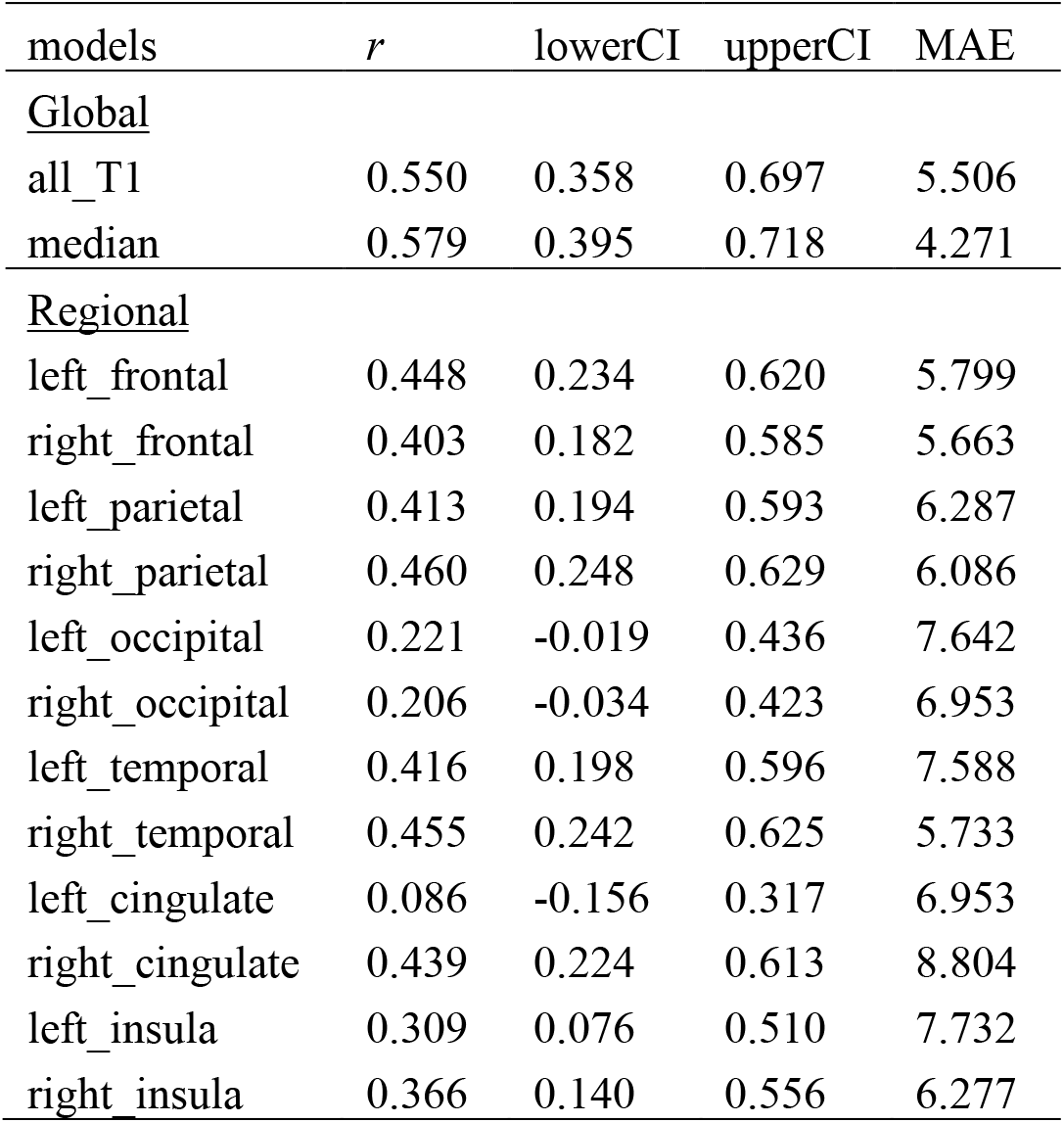
Pearson correlation between estimated brain age and chronological age with their confidence intervals on the test sample at baseline (stroke patients from StrokeMRI sample) for each model, and the MAE calculated from BAGR.

Table 3 shows ICC with their confidence intervals for each model for the two baselines and for the three timepoints ranging from .89 (CI=.82-.94) for the right parietal model to .68 (CI=.50-.80) for the left cingulate model across the two baseline assessments, and ranging from .86 (CI=.79-.91) for the right parietal model to .70 (CI=.57-.80) for the left cingulate model across the three timepoints. See Suppl. Table 2 for the estimation after replacing outliers by imputed values.

**Table 3.**
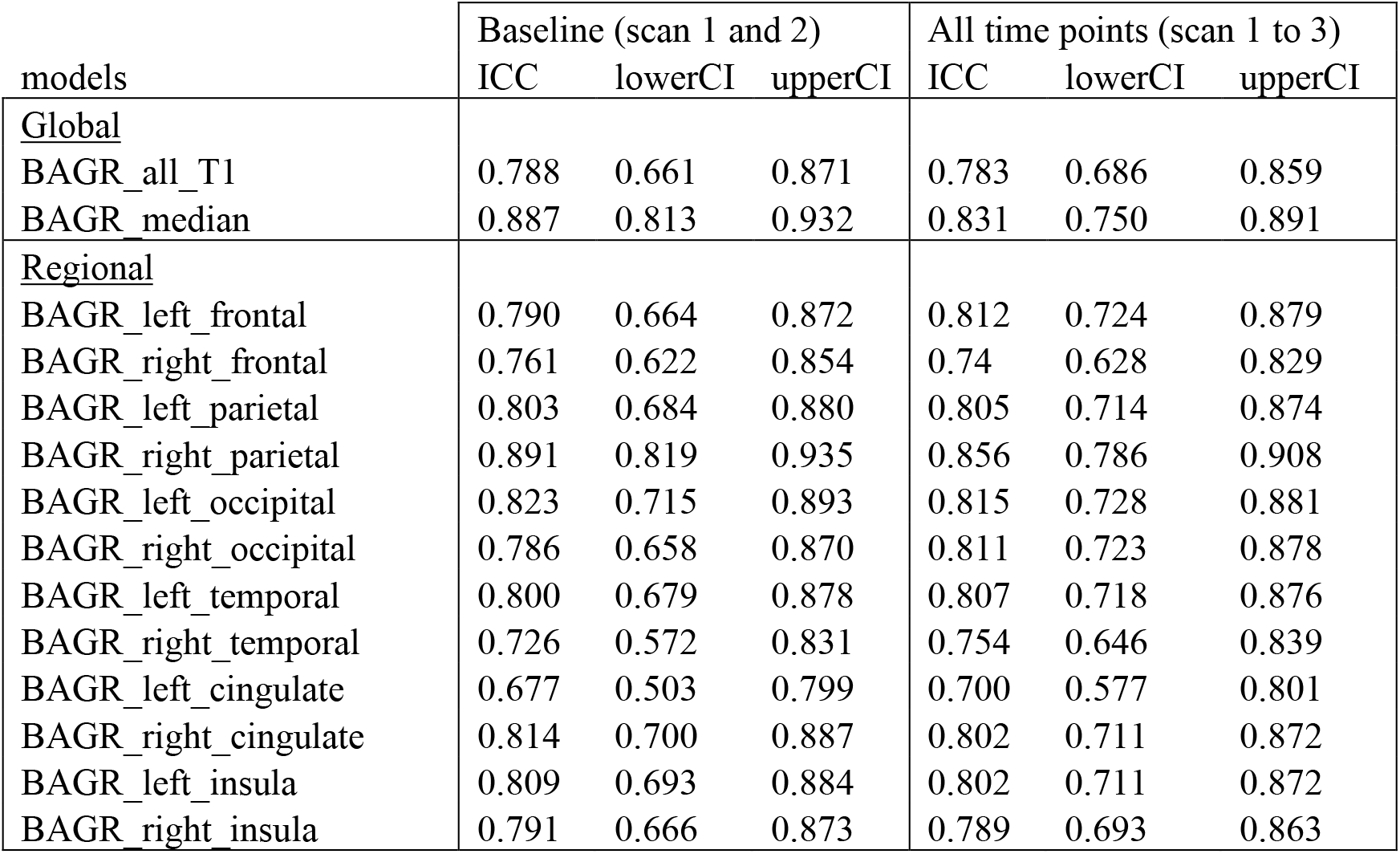
Intra-class correlation (ICC) with their confidence interval of the estimated brain age for the two baseline scans (scan one and two), and for the three timepoints (scan one, two and three).

| models | Baseline (scan 1 and 2) |  |  | All time points (scan 1 to 3) |  |  |
| --- | --- | --- | --- | --- | --- | --- |
|  | ICC | lowerCI | upperCI | ICC | lowerCI | upperCI |
| <u>Global</u> |  |  |  |  |  |  |
| BAGR_all_T1 | 0.788 | 0.661 | 0.871 | 0.783 | 0.686 | 0.859 |
| BAGR_median | 0.887 | 0.813 | 0.932 | 0.831 | 0.750 | 0.891 |
| <u>Regional</u> |  |  |  |  |  |  |
| BAGR_left_frontal | 0.790 | 0.664 | 0.872 | 0.812 | 0.724 | 0.879 |
| BAGR_right_frontal | 0.761 | 0.622 | 0.854 | 0.74 | 0.628 | 0.829 |
| BAGR_left_parietal | 0.803 | 0.684 | 0.880 | 0.805 | 0.714 | 0.874 |
| BAGR_right_parietal | 0.891 | 0.819 | 0.935 | 0.856 | 0.786 | 0.908 |
| BAGR_left_occipital | 0.823 | 0.715 | 0.893 | 0.815 | 0.728 | 0.881 |
| BAGR_right_occipital | 0.786 | 0.658 | 0.870 | 0.811 | 0.723 | 0.878 |
| BAGR_left_temporal | 0.800 | 0.679 | 0.878 | 0.807 | 0.718 | 0.876 |
| BAGR_right_temporal | 0.726 | 0.572 | 0.831 | 0.754 | 0.646 | 0.839 |
| BAGR_left_cingulate | 0.677 | 0.503 | 0.799 | 0.700 | 0.577 | 0.801 |
| BAGR_right_cingulate | 0.814 | 0.700 | 0.887 | 0.802 | 0.711 | 0.872 |
| BAGR_left_insula | 0.809 | 0.693 | 0.884 | 0.802 | 0.711 | 0.872 |
| BAGR_right_insula | 0.791 | 0.666 | 0.873 | 0.789 | 0.693 | 0.863 |

Table 4 and Table 5 show summary statistics from the linear models testing for associations between cognitive performance at baseline and Cogmed performance gain, respectively, and BAGR, including age and sex in the models. As expected, we found a main effect of age on cognitive performance at baseline. However, the analyses revealed no significant associations between cognitive performance at baseline and BAGR after FDR correction for multiple comparisons. Amongst the non-significant findings, the strongest associations were found between cluster 5 (executive control and speed) and the right cingulate, cluster 7 (working memory) and the right cingulate, and cluster 4 (attentional control and speed) and the right temporal BAGR. Further, we did not find any significant associations between performance improvement score and BAGR, nor main effect of age, nor sex on the performance improvement score after FDR corrections. Amongst the non-significant findings, the strongest associations with Cogmed performance gain were found for the left frontal and left parietal models, indicating higher cognitive gain for participants with lower BAGR.

**Table 4.** Summary statistics of the associations between cognitive performance at baseline and BAGR, including age and sex using linear models. **Cluster 1:** memory and learning. **Cluster 2:** visual processing speed. **Cluster 3:** verbal skills. **Cluster 4:** attentional control and speed. **Cluster 5:** executive control and speed. **Cluster 6:** reasoning and psychomotor speed. **Cluster 7:** working memory. The reported p-values are uncorrected values, and no main effect of BAGR remained significant after FDR correction.

| clusters | models | Bagr (t (p)) | Age (t (p)) | Sex (t (p)) |
| --- | --- | --- | --- | --- |
| Cluster1 | all_T1 | 0.152 (0.880) | -4.884 (<0.001) | -2.200 (0.032) |
| Cluster1 | median | -0.095 (0.925) | -4.886 (<0.001) | -2.220 (0.030) |
| Cluster1 | left_frontal | -0.032 (0.975) | -4.886 (<0.001) | -2.201 (0.031) |
| Cluster1 | right_frontal | -0.795 (0.43) | -4.933 (<0.001) | -2.169 (0.034) |
| Cluster1 | left_parietal | -0.356 (0.723) | -4.879 (<0.001) | -2.227 (0.030) |
| Cluster1 | right_parietal | 0.466 (0.643) | -4.902 (<0.001) | -2.187 (0.032) |
| Cluster1 | left_occipital | -0.797 (0.429) | -4.957 (<0.001) | -2.281 (0.026) |
| Cluster1 | right_occipital | 0.478 (0.634) | -4.846 (<0.001) | -2.182 (0.033) |
| Cluster1 | left_temporal | -0.283 (0.778) | -4.893 (<0.001) | -2.232 (0.029) |
| Cluster1 | right_temporal | -0.495 (0.622) | -4.906 (<0.001) | -2.246 (0.028) |
| Cluster1 | left_cingulate | 1.018 (0.313) | -4.910 (<0.001) | -2.146 (0.036) |
| Cluster1 | right_cingulate | -0.330 (0.742) | -4.843 (<0.001) | -2.230 (0.029) |
| Cluster1 | left_insula | -0.898 (0.373) | -4.921 (<0.001) | -2.322 (0.024) |
| Cluster1 | right_insula | -0.651 (0.518) | -4.922 (<0.001) | -2.176 (0.033) |
| Cluster2 | all_T1 | 0.277 (0.783) | -3.716 (<0.001) | 1.853 (0.069) |
| Cluster2 | median | 0.263 (0.793) | -3.723 (<0.001) | 1.848 (0.069) |
| Cluster2 | left_frontal | 0.120 (0.905) | -3.721 (<0.001) | 1.821 (0.073) |
| Cluster2 | right_frontal | -0.754 (0.454) | -3.753 (<0.001) | 1.884 (0.064) |
| Cluster2 | left_parietal | 0.304 (0.762) | -3.729 (<0.001) | 1.840 (0.070) |
| Cluster2 | right_parietal | -0.012 (0.991) | -3.718 (<0.001) | 1.830 (0.072) |
| Cluster2 | left_occipital | 0.122 (0.903) | -3.700 (<0.001) | 1.838 (0.071) |
| Cluster2 | right_occipital | -0.931 (0.355) | -3.806 (<0.001) | 1.782 (0.08) |
| Cluster2 | left_temporal | 1.283 (0.204) | -3.752 (<0.001) | 1.927 (0.058) |
| Cluster2 | right_temporal | 0.345 (0.731) | -3.713 (<0.001) | 1.851 (0.069) |
| Cluster2 | left_cingulate | 0.498 (0.62) | -3.716 (<0.001) | 1.870 (0.066) |
| Cluster2 | right_cingulate | -0.431 (0.668) | -3.679 (<0.001) | 1.833 (0.072) |
| Cluster2 | left_insula | -0.482 (0.632) | -3.728 (<0.001) | 1.773 (0.081) |
| Cluster2 | right_insula | -0.658 (0.513) | -3.750 (<0.001) | 1.880 (0.065) |
| Cluster3 | all_T1 | -0.497 (0.621) | -2.96 (0.004) | -3.072 (0.003) |
| Cluster3 | median | -0.529 (0.599) | -2.947 (0.004) | -3.068 (0.003) |
| Cluster3 | left_frontal | 0.986 (0.328) | -2.996 (0.004) | -3.119 (0.003) |
| Cluster3 | right_frontal | -0.744 (0.460) | -2.977 (0.004) | -2.997 (0.004) |
| Cluster3 | left_parietal | -0.808 (0.422) | -2.935 (0.005) | -3.063 (0.003) |
| Cluster3 | right_parietal | 0.796 (0.429) | -2.971 (0.004) | -2.995 (0.004) |
| Cluster3 | left_occipital | -0.608 (0.545) | -2.992 (0.004) | -3.079 (0.003) |
| Cluster3 | right_occipital | 0.243 (0.809) | -2.919 (0.005) | -3.013 (0.004) |
| Cluster3 | left_temporal | 0.264 (0.793) | -2.945 (0.004) | -3.016 (0.004) |
| Cluster3 | right_temporal | -0.901 (0.371) | -2.985 (0.004) | -3.096 (0.003) |
| Cluster3 | left_cingulate | -0.462 (0.646) | -2.959 (0.004) | -3.066 (0.003) |
| Cluster3 | right_cingulate | -1.708 (0.092) | -2.87 (0.006) | -3.118 (0.003) |
| Cluster3 | left_insula | -1.207 (0.232) | -2.987 (0.004) | -3.183 (0.002) |
| Cluster3 | right_insula | -1.909 (0.061) | -3.086 (0.003) | -2.983 (0.004) |
| Cluster4 | all_T1 | -1.687 (0.096) | -4.182 (<0.001) | -0.170 (0.866) |
| Cluster4 | median | -1.990 (0.051) | -4.166 (<0.001) | -0.142 (0.887) |
| Cluster4 | left_frontal | -0.833 (0.408) | -4.060 (<0.001) | 0.043 (0.966) |
| Cluster4 | right_frontal | -0.838 (0.405) | -4.106 (<0.001) | 0.034 (0.973) |
| Cluster4 | left_parietal | -1.325 (0.190) | -4.076 (<0.001) | -0.040 (0.968) |
| Cluster4 | right_parietal | -1.471 (0.146) | -4.112 (<0.001) | -0.106 (0.916) |
| Cluster4 | left_occipital | -1.884 (0.064) | -4.303 (<0.001) | -0.155 (0.878) |
| Cluster4 | right_occipital | -1.334 (0.187) | -4.213 (<0.001) | -0.103 (0.918) |
| Cluster4 | left_temporal | -0.280 (0.780) | -4.069 (<0.001) | -0.032 (0.974) |
| Cluster4 | right_temporal | -2.063 (0.043) | -4.243 (<0.001) | -0.125 (0.901) |
| Cluster4 | left_cingulate | 0.023 (0.982) | -4.063 (<0.001) | -0.015 (0.988) |
| Cluster4 | right_cingulate | -1.316 (0.193) | -4.003 (<0.001) | -0.029 (0.977) |
| Cluster4 | left_insula | -0.042 (0.967) | -4.064 (<0.001) | -0.021 (0.983) |
| Cluster4 | right insula | -0.637 (0.527) | -4.094 (<0.001) | 0.026 (0.979) |
| Cluster5 | all_T1 | -1.318 (0.192) | -4.442 (<0.001) | -2.098 (0.040) |
| Cluster5 | median | -1.427 (0.158) | -4.415 (<0.001) | -2.077 (0.042) |
| Cluster5 | left_frontal | 0.570 (0.571) | -4.388 (<0.001) | -2.000 (0.050) |
| Cluster5 | right_frontal | -0.82 (0.415) | -4.404 (<0.001) | -1.917 (0.060) |
| Cluster5 | left_parietal | 0.464 (0.644) | -4.381 (<0.001) | -1.955 (0.055) |
| Cluster5 | right_parietal | -0.449 (0.655) | -4.362 (<0.001) | -1.986 (0.051) |
| Cluster5 | left_occipital | -0.417 (0.678) | -4.386 (<0.001) | -1.988 (0.051) |
| Cluster5 | right_occipital | -1.995 (0.050) | -4.639 (<0.001) | -2.145 (0.036) |
| Cluster5 | left_temporal | -1.263 (0.211) | -4.430 (<0.001) | -2.052 (0.044) |
| Cluster5 | right_temporal | -1.827 (0.072) | -4.515 (<0.001) | -2.103 (0.039) |
| Cluster5 | left_cingulate | 1.429 (0.158) | -4.405 (<0.001) | -1.874 (0.066) |
| Cluster5 | right_cingulate | -2.387 (0.020) | -4.351 (<0.001) | -2.067 (0.043) |
| Cluster5 | left_insula | -1.840 (0.070) | -4.487 (<0.001) | -2.202 (0.031) |
| Cluster5 | right insula | 0.655 (0.515) | -4.354 (<0.001) | -2.006 (0.049) |
| Cluster6 | all_T1 | -0.008 (0.993) | -5.334 (<0.001) | -2.252 (0.028) |
| Cluster6 | median | -0.278 (0.782) | -5.334 (<0.001) | -2.275 (0.026) |
| Cluster6 | left_frontal | 0.176 (0.860) | -5.339 (<0.001) | -2.268 (0.027) |
| Cluster6 | right_frontal | -1.132 (0.262) | -5.415 (<0.001) | -2.211 (0.031) |
| Cluster6 | left_parietal | -0.128 (0.898) | -5.329 (<0.001) | -2.263 (0.027) |
| Cluster6 | right_parietal | 0.309 (0.758) | -5.342 (<0.001) | -2.239 (0.029) |
| Cluster6 | left_occipital | -1.046 (0.300) | -5.443 (<0.001) | -2.350 (0.022) |
| Cluster6 | right_occipital | -0.179 (0.858) | -5.333 (<0.001) | -2.268 (0.027) |
| Cluster6 | left_temporal | 0.673 (0.503) | -5.346 (<0.001) | -2.227 (0.030) |
| Cluster6 | right_temporal | -0.280 (0.781) | -5.343 (<0.001) | -2.273 (0.026) |
| Cluster6 | left_cingulate | 0.825 (0.412) | -5.347 (<0.001) | -2.202 (0.031) |
| Cluster6 | right_cingulate | -1.443 (0.154) | -5.291 (<0.001) | -2.310 (0.024) |
| Cluster6 | left_insula | -1.027 (0.308) | -5.385 (<0.001) | -2.378 (0.020) |
| Cluster6 | right insula | 0.013 (0.990) | -5.332 (<0.001) | -2.256 (0.028) |
| Cluster7 | all T1 | 0.206 (0.838) | -5.298 (<0.001) | -1.473 (0.146) |
| Cluster7 | median | -0.431 (0.668) | -5.303 (<0.001) | -1.524 (0.132) |
| Cluster7 | left_frontal | 0.636 (0.527) | -5.333 (<0.001) | -1.544 (0.128) |
| Cluster7 | right_frontal | 0.003 (0.998) | -5.298 (<0.001) | -1.495 (0.140) |
| Cluster7 | left_parietal | -0.378 (0.707) | -5.292 (<0.001) | -1.506 (0.137) |
| Cluster7 | right_parietal | 0.151 (0.881) | -5.303 (<0.001) | -1.486 (0.142) |
| Cluster7 | left_occipital | -0.595 (0.554) | -5.344 (<0.001) | -1.541 (0.128) |
| Cluster7 | right_occipital | -1.219 (0.227) | -5.441 (<0.001) | -1.591 (0.117) |
| Cluster7 | left_temporal | 0.799 (0.427) | -5.317 (<0.001) | -1.458 (0.150) |
| Cluster7 | right_temporal | -0.336 (0.738) | -5.311 (<0.001) | -1.515 (0.135) |
| Cluster7 | left_cingulate | -0.572 (0.569) | -5.323 (<0.001) | -1.541 (0.128) |
| Cluster7 | right_cingulate | -2.241 (0.028) | -5.312 (<0.001) | -1.576 (0.120) |
| Cluster7 | left_insula | 0.989 (0.326) | -5.335 (<0.001) | -1.391 (0.169) |
| Cluster7 | right_insula | -0.234 (0.816) | -5.307 (<0.001) | -1.479 (0.144) |

**Table 5.** Summary statistics of the associations between Cogmed performance gain and BAGR, including age and sex using linear models. The reported p-values are uncorrected values, and no main effect of BAGR remained significant after FDR correction.

| models | Bagr (t (p)) | Age (t (p)) | Sex (t (p)) |
| --- | --- | --- | --- |
| <u>Global</u> |  |  |  |
| BAGR_all_T1 | -0.383 (0.703) | 0.497 (0.622) | -0.900 (0.373) |
| BAGR_median | 0.382 (0.704) | 0.453 (0.652) | -0.902 (0.371) |
| <u>Regional</u> |  |  |  |
| BAGR_left_frontal | 2.399 (0.020) | 0.285 (0.777) | -1.146 (0.257) |
| BAGR_right_frontal | 0.241 (0.810) | 0.479 (0.634) | -0.913 (0.366) |
| BAGR_left_parietal | 2.037 (0.047) | 0.425 (0.673) | -0.876 (0.385) |
| BAGR_right_parietal | 0.264 (0.793) | 0.493 (0.625) | -0.899 (0.373) |
| BAGR_left_occipital | -0.083 (0.934) | 0.489 (0.627) | -0.899 (0.373) |
| BAGR_right_occipital | 0.392 (0.697) | 0.502 (0.618) | -0.909 (0.368) |
| BAGR_left_temporal | 0.159 (0.875) | 0.484 (0.631) | -0.900 (0.372) |
| BAGR_right_temporal | 0.963 (0.340) | 0.452 (0.653) | -0.925 (0.359) |
| BAGR_left_cingulate | 0.750 (0.457) | 0.438 (0.663) | -0.883 (0.381) |
| BAGR_right_cingulate | 1.150 (0.256) | 0.436 (0.664) | -1.053 (0.297) |
| BAGR_left_insula | 0.458 (0.649) | 0.487 (0.629) | -0.882 (0.382) |
| BAGR_right_insula | 0.693 (0.491) | 0.452 (0.653) | -0.952 (0.345) |

Table 6 shows summary statistics from the linear mixed effects models testing for longitudinal associations between cognitive performance and BAGR, including age and sex in the models. The analyses revealed robust main effects of session and age, indicating increasing performance during the course of the intervention, and generally lower performance with increasing age. Beyond this, we did not find any significant associations between performance and BAGR, nor BAGR by time interaction after FDR correction for multiple comparisons. Amongst the non-significant findings, the five strongest associations were found between the left occipital and *Digits*, the left frontal and *Cube* and *3D Cube*, the left parietal and *Hidden*, and the right insula and *Twist* and the eight strongest BAGR by time interactions were found between the left occipital and *Digits*, the left frontal and *Cube* and *3D Cube*, the left parietal and *Hidden*, the right insula and *Twist* and *Rotating*, and the left temporal and *Cube* and *3D Cube*.

**Table 6.** Summary statistics from the linear mixed effects models testing for associations between cognitive performance and BAGR by time interaction, including age and sex in the models. The reported p-values are uncorrected values, and no main effect of BAGR nor BAGR by time interaction remained significant after FDR correction.

| test | models | Bagr (t (p)) | Session (t (p)) | Age (t (p)) | Sex (t (p)) | bagr:session (t (p)) |
| --- | --- | --- | --- | --- | --- | --- |
| Grid | BAGR_all_T1 | -0.478 (0.635) | 10.704 (<0.001) | -3.538 (0.001) | -1.064 (0.292) | 0.338 (0.737) |
| Sort | BAGR_all_T1 | -1.409 (0.165) | 10.097 (<0.001) | -2.642 (0.011) | -0.573 (0.569) | 1.157 (0.253) |
| Digits | BAGR_all_T1 | 0.739 (0.463) | 10.198 (<0.001) | -6.649 (<0.001) | -2.507 (0.015) | -0.685 (0.496) |
| Cube | BAGR_all_T1 | 0.053 (0.958) | 15.279 (<0.001) | -3.047 (0.004) | -1.384 (0.172) | -0.211 (0.833) |
| Hidden | BAGR_all_T1 | -0.213 (0.832) | 9.487 (<0.001) | -5.612 (<0.001) | -2.764 (0.008) | 0.929 (0.357) |
| Twist | BAGR_all_T1 | -0.230 (0.819) | 6.202 (<0.001) | -3.813 (<0.001) | -1.453 (0.152) | 0.088 (0.930) |
| 3D Cube | BAGR_all_T1 | 1.444 (0.155) | 6.369 (<0.001) | -4.353 (<0.001) | -1.574 (0.122) | -1.631 (0.109) |
| Rotating | BAGR_all_T1 | 0.198 (0.844) | 4.307 (<0.001) | -3.984 (<0.001) | -1.508 (0.138) | -0.349 (0.729) |
| Grid | BAGR_median | -0.507 (0.615) | 10.840 (<0.001) | -3.521 (0.001) | -1.050 (0.299) | 0.116 (0.908) |
| Sort | BAGR_median | -1.288 (0.204) | 10.063 (<0.001) | -2.648 (0.011) | -0.610 (0.544) | 0.968 (0.338) |
| Digits | BAGR_median | 1.091 (0.280) | 10.377 (<0.001) | -6.590 (<0.001) | -2.472 (0.017) | -1.165 (0.250) |
| Cube | BAGR_median | 0.157 (0.876) | 15.422 (<0.001) | -3.043 (0.004) | -1.376 (0.175) | -0.533 (0.596) |
| Hidden | BAGR_median | 0.357 (0.722) | 9.768 (<0.001) | -5.420 (<0.001) | -2.665 (0.010) | 0.286 (0.776) |
| Twist | BAGR_median | 0.520 (0.605) | 6.203 (<0.001) | -3.816 (<0.001) | -1.505 (0.138) | -0.614 (0.542) |
| 3D Cube | BAGR_median | 1.345 (0.185) | 6.454 (<0.001) | -4.335 (<0.001) | -1.522 (0.134) | -1.808 (0.077) |
| Rotating | BAGR_median | 0.894 (0.376) | 4.327 (<0.001) | -3.982 (<0.001) | -1.538 (0.130) | -1.072 (0.289) |
| Grid | BAGR_left_frontal | 0.298 (0.767) | 10.94 (<0.001) | -3.574 (0.001) | -1.165 (0.249) | -0.981 (0.331) |
| Sort | BAGR_left_frontal | -0.020 (0.984) | 10.125 (<0.001) | -2.661 (0.011) | -0.749 (0.457) | -0.624 (0.536) |
| Digits | BAGR_left_frontal | 1.800 (0.078) | 10.527 (<0.001) | -6.643 (<0.001) | -2.450 (0.018) | -1.942 (0.058) |
| Cube | BAGR_left_frontal | 2.423 (0.019) | 16.704 (<0.001) | -3.086 (0.003) | -1.439 (0.156) | -3.187 (0.003) |
| Hidden | BAGR_left_frontal | 1.343 (0.186) | 9.835 (<0.001) | -5.277 (<0.001) | -2.502 (0.016) | -1.278 (0.207) |
| Twist | BAGR_left_frontal | 1.333 (0.189) | 6.316 (<0.001) | -3.850 (<0.001) | -1.524 (0.134) | -1.667 (0.102) |
| 3D Cube | BAGR_left_frontal | 2.274 (0.027) | 6.407 (<0.001) | -4.419 (<0.001) | -1.537 (0.130) | -2.380 (0.021) |
| Rotating | BAGR_left_frontal | 1.496 (0.141) | 4.297 (<0.001) | -4.025 (<0.001) | -1.529 (0.132) | -1.526 (0.133) |
| Grid | BAGR_right_frontal | 0.024 (0.981) | 10.660 (<0.001) | -3.554 (0.001) | -1.117 (0.269) | -0.019 (0.985) |
| Sort | BAGR_right_frontal | 0.417 (0.679) | 10.034 (<0.001) | -2.631 (0.011) | -0.681 (0.499) | -0.187 (0.853) |
| Digits | BAGR_right_frontal | -0.428 (0.671) | 10.187 (<0.001) | -6.666 (<0.001) | -2.475 (0.017) | 0.324 (0.747) |
| Cube | BAGR_right_frontal | 1.340 (0.186) | 15.643 (<0.001) | -3.027 (0.004) | -1.387 (0.171) | -1.412 (0.164) |
| Hidden | BAGR_right_frontal | -0.501 (0.618) | 9.721 (<0.001) | -5.308 (<0.001) | -2.535 (0.014) | 0.592 (0.556) |
| Twist | BAGR_right_frontal | 0.012 (0.990) | 6.264 (<0.001) | -3.789 (<0.001) | -1.503 (0.139) | -0.320 (0.750) |
| 3D Cube | BAGR_right_frontal | 1.279 (0.207) | 6.294 (<0.001) | -4.351 (<0.001) | -1.513 (0.136) | -1.269 (0.210) |
| Rotating | BAGR_right_frontal | 0.330 (0.743) | 4.298 (<0.001) | -3.980 (<0.001) | -1.526 (0.133) | -0.464 (0.645) |
| Grid | BAGR_left_parietal | 0.879 (0.384) | 10.990 (<0.001) | -3.536 (0.001) | -1.094 (0.279) | -1.331 (0.189) |
| Sort | BAGR_left_parietal | -0.197 (0.845) | 10.109 (<0.001) | -2.665 (0.010) | -0.661 (0.511) | -0.344 (0.733) |
| Digits | BAGR_left_parietal | 0.968 (0.338) | 10.668 (<0.001) | -6.620 (<0.001) | -2.417 (0.019) | -1.482 (0.145) |
| Cube | BAGR_left_parietal | 0.075 (0.941) | 15.499 (<0.001) | -3.044 (0.004) | -1.377 (0.174) | -0.514 (0.610) |
| Hidden | BAGR_left_parietal | 2.051 (0.046) | 10.062 (<0.001) | -5.262 (<0.001) | -2.544 (0.014) | -2.060 (0.045) |
| Twist | BAGR_left_parietal | 1.535 (0.131) | 6.406 (<0.001) | -3.804 (<0.001) | -1.496 (0.141) | -1.775 (0.082) |
| 3D Cube | BAGR_left_parietal | 0.291 (0.772) | 6.561 (<0.001) | -4.398 (<0.001) | -1.493 (0.141) | -1.054 (0.297) |
| Rotating | BAGR_left_parietal | 1.225 (0.227) | 4.423 (<0.001) | -3.986 (<0.001) | -1.521 (0.134) | -1.515 (0.136) |
| Grid | BAGR_right_parietal | -0.990 (0.327) | 10.795 (<0.001) | -3.558 (0.001) | -1.072 (0.289) | 0.661 (0.512) |
| Sort | BAGR_right_parietal | -0.489 (0.627) | 9.883 (<0.001) | -2.661 (0.010) | -0.673 (0.504) | 0.248 (0.805) |
| Digits | BAGR_right_parietal | 0.202 (0.841) | 10.461 (<0.001) | -6.620 (<0.001) | -2.432 (0.019) | -0.712 (0.480) |
| Cube | BAGR_right_parietal | -0.327 (0.745) | 15.394 (<0.001) | -3.058 (0.004) | -1.385 (0.172) | -0.116 (0.908) |
| Hidden | BAGR_right_parietal | -0.450 (0.655) | 9.734 (<0.001) | -5.344 (<0.001) | -2.544 (0.014) | 0.817 (0.418) |
| Twist | BAGR_right_parietal | -0.226 (0.822) | 6.180 (<0.001) | -3.832 (<0.001) | -1.487 (0.143) | 0.086 (0.932) |
| 3D Cube | BAGR_right_parietal | 0.994 (0.325) | 6.366 (<0.001) | -4.386 (<0.001) | -1.554 (0.126) | -1.489 (0.143) |
| Rotating | BAGR_right_parietal | 0.625 (0.535) | 4.309 (<0.001) | -3.984 (<0.001) | -1.543 (0.129) | -0.716 (0.477) |
| Grid | BAGR_left_occipital | -1.717 (0.092) | 11.041 (<0.001) | -3.487 (0.001) | -1.095 (0.279) | 1.794 (0.079) |
| Sort | BAGR_left_occipital | -0.489 (0.627) | 10.050 (<0.001) | -2.561 (0.014) | -0.664 (0.510) | 0.282 (0.779) |
| Digits | BAGR_left_occipital | 2.399 (0.020) | 10.716 (<0.001) | -6.684 (<0.001) | -2.480 (0.016) | -2.383 (0.021) |
| Cube | BAGR_left_occipital | -0.202 (0.840) | 15.254 (<0.001) | -3.083 (0.003) | -1.423 (0.161) | 0.473 (0.638) |
| Hidden | BAGR_left_occipital | 0.028 (0.978) | 9.630 (<0.001) | -5.413 (<0.001) | -2.599 (0.012) | 0.407 (0.685) |
| Twist | BAGR_left_occipital | -0.265 (0.792) | 6.197 (<0.001) | -3.782 (<0.001) | -1.483 (0.144) | 0.144 (0.886) |
| 3D Cube | BAGR_left_occipital | 0.340 (0.735) | 6.246 (<0.001) | -4.33 (<0.001) | -1.515 (0.136) | -0.510 (0.612) |
| Rotating | BAGR_left_occipital | -0.718 (0.476) | 4.271 (<0.001) | -4.005 (<0.001) | -1.54 (0.130) | 0.899 (0.373) |
| Grid | BAGR_right_occipital | -0.632 (0.530) | 10.557 (<0.001) | -3.532 (0.001) | -1.061 (0.294) | 0.236 (0.814) |
| Sort | BAGR_right_occipital | -1.314 (0.195) | 9.815 (<0.001) | -2.613 (0.012) | -0.621 (0.537) | 1.026 (0.310) |
| Digits | BAGR_right_occipital | 0.595 (0.555) | 10.645 (<0.001) | -6.638 (<0.001) | -2.400 (0.020) | -1.355 (0.182) |
| Cube | BAGR_right_occipital | -0.982 (0.331) | 15.919 (<0.001) | -2.993 (0.004) | -1.297 (0.200) | -0.201 (0.842) |
| Hidden | BAGR_right_occipital | -0.265 (0.792) | 9.655 (<0.001) | -5.254 (<0.001) | -2.478 (0.016) | -0.063 (0.950) |
| Twist | BAGR_right_occipital | -0.578 (0.566) | 6.114 (<0.001) | -3.808 (<0.001) | -1.433 (0.158) | 0.065 (0.948) |
| 3D Cube | BAGR_right_occipital | 0.311 (0.757) | 6.414 (<0.001) | -4.349 (<0.001) | -1.470 (0.148) | -1.135 (0.262) |
| Rotating | BAGR_right_occipital | -0.448 (0.656) | 4.202 (<0.001) | -4.007 (<0.001) | -1.480 (0.145) | -0.052 (0.959) |
| Grid | BAGR_left_temporal | 0.192 (0.849) | 10.683 (<0.001) | -3.572 (0.001) | -1.073 (0.288) | -0.517 (0.608) |
| Sort | BAGR_left_temporal | -0.300 (0.766) | 9.878 (<0.001) | -2.656 (0.011) | -0.687 (0.495) | 0.189 (0.851) |
| Digits | BAGR_left_temporal | 0.733 (0.467) | 10.26 (<0.001) | -6.645 (<0.001) | -2.469 (0.017) | -0.552 (0.583) |
| Cube | BAGR_left_temporal | 1.224 (0.227) | 16.324 (<0.001) | -3.078 (0.003) | -1.293 (0.202) | -2.241 (0.030) |
| Hidden | BAGR_left_temporal | 0.367 (0.715) | 9.897 (<0.001) | -5.359 (<0.001) | -2.665 (0.010) | 0.33 (0.743) |
| Twist | BAGR_left_temporal | 0.160 (0.874) | 6.176 (<0.001) | -3.829 (<0.001) | -1.492 (0.142) | -0.18 (0.858) |
| 3D Cube | BAGR_left_temporal | 1.516 (0.136) | 6.551 (<0.001) | -4.407 (<0.001) | -1.412 (0.164) | -2.271 (0.028) |
| Rotating | BAGR_left_temporal | 0.480 (0.633) | 4.297 (<0.001) | -3.988 (<0.001) | -1.527 (0.133) | -0.403 (0.689) |
| Grid | BAGR_right_temporal | 0.116 (0.908) | 10.638 (<0.001) | -3.552 (0.001) | -1.120 (0.268) | -0.128 (0.899) |
| Sort | BAGR_right_temporal | -0.530 (0.599) | 9.997 (<0.001) | -2.660 (0.011) | -0.679 (0.500) | 0.684 (0.497) |
| Digits | BAGR_right_temporal | 1.236 (0.222) | 10.329 (<0.001) | -6.602 (<0.001) | -2.513 (0.015) | -1.445 (0.155) |
| Cube | BAGR_right_temporal | 0.791 (0.433) | 15.444 (<0.001) | -3.033 (0.004) | -1.417 (0.162) | -1.110 (0.272) |
| Hidden | BAGR_right_temporal | 1.014 (0.315) | 9.635 (<0.001) | -5.331 (<0.001) | -2.620 (0.011) | -0.902 (0.372) |
| Twist | BAGR_right_temporal | -0.183 (0.856) | 6.233 (<0.001) | -3.789 (<0.001) | -1.456 (0.151) | -0.011 (0.991) |
| 3D Cube | BAGR_right_temporal | 1.417 (0.163) | 6.186 (<0.001) | -4.397 (<0.001) | -1.630 (0.109) | -1.332 (0.189) |
| Rotating | BAGR_right_temporal | -0.168 (0.867) | 4.288 (<0.001) | -3.991 (<0.001) | -1.506 (0.138) | 0.077 (0.939) |
| Grid | BAGR_left_cingulate | 0.182 (0.856) | 10.635 (<0.001) | -3.531 (0.001) | -1.076 (0.287) | -0.369 (0.714) |
| Sort | BAGR_left_cingulate | 0.667 (0.508) | 9.662 (<0.001) | -2.683 (0.010) | -0.746 (0.459) | -0.481 (0.633) |
| Digits | BAGR_left_cingulate | 0.318 (0.752) | 10.09 (<0.001) | -6.641 (<0.001) | -2.440 (0.018) | -0.345 (0.731) |
| Cube | BAGR_left_cingulate | -1.008 (0.318) | 15.291 (<0.001) | -3.076 (0.003) | -1.441 (0.156) | 1.259 (0.214) |
| Hidden | BAGR_left_cingulate | -0.726 (0.472) | 9.718 (<0.001) | -5.376 (<0.001) | -2.661 (0.010) | 1.235 (0.223) |
| Twist | BAGR_left_cingulate | -0.550 (0.585) | 6.115 (<0.001) | -3.860 (<0.001) | -1.546 (0.128) | 0.737 (0.465) |
| 3D Cube | BAGR_left_cingulate | 0.251 (0.803) | 6.225 (<0.001) | -4.348 (<0.001) | -1.474 (0.146) | -0.504 (0.617) |
| Rotating | BAGR_left_cingulate | 0.524 (0.603) | 4.161 (<0.001) | -4.013 (<0.001) | -1.546 (0.128) | -0.453 (0.652) |
| Grid | BAGR_right_cingulate | -0.486 (0.629) | 10.575 (<0.001) | -3.564 (0.001) | -1.112 (0.271) | 0.353 (0.725) |
| Sort | BAGR_right_cingulate | 0.034 (0.973) | 9.922 (<0.001) | -2.625 (0.012) | -0.706 (0.484) | 0.099 (0.921) |
| Digits | BAGR_right_cingulate | 0.266 (0.792) | 10.123 (<0.001) | -6.658 (<0.001) | -2.447 (0.018) | -0.333 (0.741) |
| Cube | BAGR_right_cingulate | -0.263 (0.794) | 15.110 (<0.001) | -3.116 (0.003) | -1.39 (0.171) | -0.248 (0.805) |
| Hidden | BAGR_right_cingulate | 0.390 (0.698) | 9.694 (<0.001) | -5.264 (<0.001) | -2.506 (0.015) | -0.424 (0.674) |
| Twist | BAGR_right_cingulate | 0.684 (0.497) | 6.138 (<0.001) | -3.846 (<0.001) | -1.493 (0.142) | -0.737 (0.465) |
| 3D Cube | BAGR_right_cingulate | 1.072 (0.289) | 6.119 (<0.001) | -4.48 (<0.001) | -1.511 (0.137) | -1.521 (0.135) |
| Rotating | BAGR_right_cingulate | 0.262 (0.794) | 4.352 (<0.001) | -3.955 (<0.001) | -1.533 (0.131) | -0.056 (0.956) |
| Grid | BAGR_left_insula | 0.704 (0.485) | 10.678 (<0.001) | -3.568 (0.001) | -1.153 (0.254) | -0.543 (0.590) |
| Sort | BAGR_left_insula | -1.289 (0.204) | 10.083 (<0.001) | -2.639 (0.011) | -0.663 (0.510) | 1.223 (0.227) |
| Digits | BAGR_left_insula | 1.611 (0.114) | 10.372 (<0.001) | -6.716 (<0.001) | -2.564 (0.013) | -1.274 (0.209) |
| Cube | BAGR_left_insula | 0.282 (0.779) | 15.283 (<0.001) | -3.064 (0.004) | -1.422 (0.161) | -0.174 (0.863) |
| Hidden | BAGR_left_insula | 0.406 (0.686) | 9.749 (<0.001) | -5.526 (<0.001) | -2.753 (0.008) | 0.280 (0.781) |
| Twist | BAGR_left_insula | 0.570 (0.571) | 6.193 (<0.001) | -3.848 (<0.001) | -1.555 (0.126) | -0.323 (0.748) |
| 3D Cube | BAGR_left_insula | 1.625 (0.111) | 6.310 (<0.001) | -4.350 (<0.001) | -1.510 (0.137) | -1.862 (0.069) |
| Rotating | BAGR_left_insula | 1.102 (0.276) | 4.277 (<0.001) | -3.982 (<0.001) | -1.530 (0.132) | -1.143 (0.258) |
| Grid | BAGR_right_insula | -0.205 (0.838) | 10.677 (<0.001) | -3.546 (0.001) | -1.107 (0.273) | 0.138 (0.890) |
| Sort | BAGR_right_insula | -0.291 (0.772) | 9.912 (<0.001) | -2.637 (0.011) | -0.687 (0.495) | 0.156 (0.877) |
| Digits | BAGR_right_insula | -0.749 (0.457) | 10.191 (<0.001) | -6.797 (<0.001) | -2.417 (0.019) | 1.071 (0.289) |
| Cube | BAGR_right_insula | -0.928 (0.358) | 15.415 (<0.001) | -3.053 (0.004) | -1.372 (0.176) | 0.779 (0.440) |
| Hidden | BAGR_right_insula | 0.486 (0.629) | 9.671 (<0.001) | -5.345 (<0.001) | -2.560 (0.013) | -0.252 (0.802) |
| Twist | BAGR_right_insula | 2.631 (0.011) | 6.561 (<0.001) | -3.843 (<0.001) | -1.656 (0.104) | -2.528 (0.015) |
| 3D Cube | BAGR_right_insula | -0.859 (0.394) | 6.227 (<0.001) | -4.392 (<0.001) | -1.486 (0.143) | 0.889 (0.378) |
| Rotating | BAGR_right_insula | 1.919 (0.061) | 4.470 (<0.001) | -3.970 (<0.001) | -1.635 (0.108) | -2.041 (0.047) |

## Discussion

Cognitive deficits are important predictors for outcome, independence and quality of life in stroke survivors, and computerized cognitive training has been suggested among the candidate interventions that may alleviate them. However, the lack of widely adapted tools for stratification, outcome prediction and treatment monitoring prevent an adequate assessment of the effectiveness of such training. Advanced brain MRI provides various candidate markers for disease monitoring and outcome prediction, integrating lesion specific information and characterization of the integrity of the unaffected parts of the brain, which is highly relevant for cognitive functions and long-term outcome. Here, we used brain age prediction based on brain morphometry and machine learning to test the hypotheses that patients with younger-looking brains would show preserved cognitive function compared to patients with older-looking brains and show more beneficial treatment response.

Based on the notion that brain age prediction offers a sensitive summary measure of brain health and integrity, we first tested the prediction accuracy and then the reliability in a longitudinal context. The estimated performance of the 13 trained models within the training set using a 10-fold cross-validation procedure suggested a relatively good model fit, with correlations ranging from .84 to .61 for the model based on all T1 features and for the right cingulate model respectively. Further, the models estimated brain age on the test sample also suggested an acceptable model fit with some regional differences in performance with MAE ranging from 5.50 to 8.80 for the most comprehensive model based on all T1 features and for the model based on right cingulate features respectively. In addition, the age estimation based on the median of the 12 regional models achieved the highest performance with a Pearson correlation of *r* = .58 (CI=.40-.72, MAE=4.27).

In line with a recent implementation in patients with MS (Høgestøl et al. 2019a), our results demonstrated high reliability across all timepoints for the global and regional models with ICC ranging from .70 to .86 for the right occipital and the right parietal models respectively. The brain age estimation based on the median of the 12 regional model was amongst the most reliable models with an ICC of .89. across the two baselines and .83 across all timepoints, outperforming the estimation based on all T1 features.

A particular challenge in clinical neuroimaging is that lesions may interfere with tools for automated processing and brain segmentations. Here, to test for the influence of brain lesions on BAGR estimation and reliability, we used outlier detection on the individual features level to identify extreme observations and replace those extremes by means of imputation using predictive mean matching. Comparisons between brain age from the raw and imputed feature sets revealed only minimal influence of outliers on both brain age estimates and reliability. Importantly, this suggests that our model predictions are robust to gross segmentation errors caused by the lesions, which supports the feasibility of automated brain age prediction in patient groups with brain disorders and lesions.

To test the hypotheses that patients with low BAGR at baseline show better cognitive function and a more positive treatment response, we used seven summary scores derived from a set of neuropsychological and computerized tests assumed to be sensitive to cognitive aging at baseline and the performance gain during the course of the intensive training period respectively. Contrary to our hypothesis, after corrections for multiple comparisons, linear models revealed no significant associations between BAGR and summary scores from the baseline assessment or performance gain. In line with a few previous studies (Boyle et al. 2019; Høgestøl et al. 2019b), region specific models revealed putative associations between summary scores for executive control and speed, working memory and the right cingulate BAGR, as well as, attentional control and speed and the right temporal BAGR, suggesting better cognitive performance with lower BAGR. In addition, region specific analysis revealed non-significant putative associations between performance gain and left frontal and left parietal BAGR, indicating more positive treatment response for patients with lower BAGR. Although future studies are needed to confirm the results, these preliminary non-significant findings provide some support to the general notion that individuals with younger-appearing brains, which may reflect relevant aspects of brain health and reserve, both perform better on cognitive tests, and respond more positively to cognitive interventions.

In order to assess if cognitive gains in response to intensive cognitive training are reflected in longitudinal changes in brain age during the course of the intervention, we tested for interactions between BAGR and session on cognitive performance using linear mixed effects models. Our analyses revealed a few putative associations between BAGR and Cogmed performance, and also session by BAGR interactions on cognitive performance in a longitudinal context. However, none of these associations and interactions remained significant after correction for multiple comparisons, and the direction of the associations, if any, did not seem to converge. Hence, our results did not provide significant support for our hypothesis that cognitive gains would be reflected in longitudinal changes in brain age during the course of the intervention.

While this study does not provide significant support for the utility of brain age estimation as a sensitive measure for cognitive reserve and potential predictor for training outcomes in stroke patients, our results suggest that region specific brain age estimations are not only reliable measures, but might also be more informative than global brain age as a measure of brain health. Future studies are needed to confirm the putative associations between summary scores for executive control and speed (cluster 5) and working memory (cluster 7) and the right cingulate BAGR, as well as summary scores for attentional control and speed (cluster 4) and the right temporal BAGR; in addition to the weak associations found between the left frontal and left parietal brain age with the rate of cognitive improvement. In general, these findings are in line with our recent study suggesting that, by capturing distinct measures of brain aging, tissue specific age prediction models might better inform us about the individual determinants and heterogeneity of the aging brain compared to models collapsing several brain compartments (Richard et al. 2018).

The following methodological considerations should be taken into account while interpreting the current results. Although our patient sample was highly heterogenous in terms of location, extent of the lesions and stroke etiologies, most of them had small lesions. Further studies are needed to confirm that the current approach for brain age prediction based on automated brain morphometry is also feasible for patients with larger lesions, and how stroke etiologies can impact BAGR and cognitive improvement, for instance progressive vascular disease (small vessel and large vessel disease) may affect cognition prior to the stroke (Ihle-Hansen et al. 2014). In addition, the patients included in this study suffered from moderate to mild stroke (NIHSS < 7 at hospital discharge), representing a high functioning group with mild cognitive deficits and better overall prognosis, limiting the generalizability of our findings. Although previous studies have reported links between cognitive function and brain age in healthy controls, it is conceivable that the current associations would be stronger had we sampled from a wider distribution in terms of stroke severity and cognitive symptoms. The lack of control group prevents us from distinguishing between the time-related changes and training-related changes (Kolskår et al. 2019), and, although tailored to the current cognitive intervention regime, a longer interval between time points might have increased sensitivity to detect relevant associations between cognitive changes and changes in brain age.

In conclusion, reliable and non-invasive markers of brain health and cognitive function are needed to help improving the current treatment programs targeting cognitive deficits following stroke and other brain disorders. Although our current results do not support an immediate clinical utility of brain age prediction in highly functioning stroke patients, we speculate that using region specific brain age model as opposed to reducing the whole brain to one summary score might be as reliable and more informative potential biomarkers of brain integrity and health. Importantly, our study supports the feasibility of automated brain age prediction in patient groups with brain disorders and lesions by highlighting the minimal impact of lesions on brain age estimations.

## Acknowledgements and funding

This study was supported by the Norwegian ExtraFoundation for Health and Rehabilitation (2015/FO5146), the Research Council of Norway (249795, 248238), the South-Eastern Norway Regional Health Authority (2014097, 2015044, 2015073), Sunnaas Rehabilitation Hospital, and the Department of Psychology, University of Oslo.

