## Supplementary Fig. 1 for "Reliable longitudinal brain age prediction in stroke patients: Associations with cognitive function and response to cognitive training"

Supplementary Figures

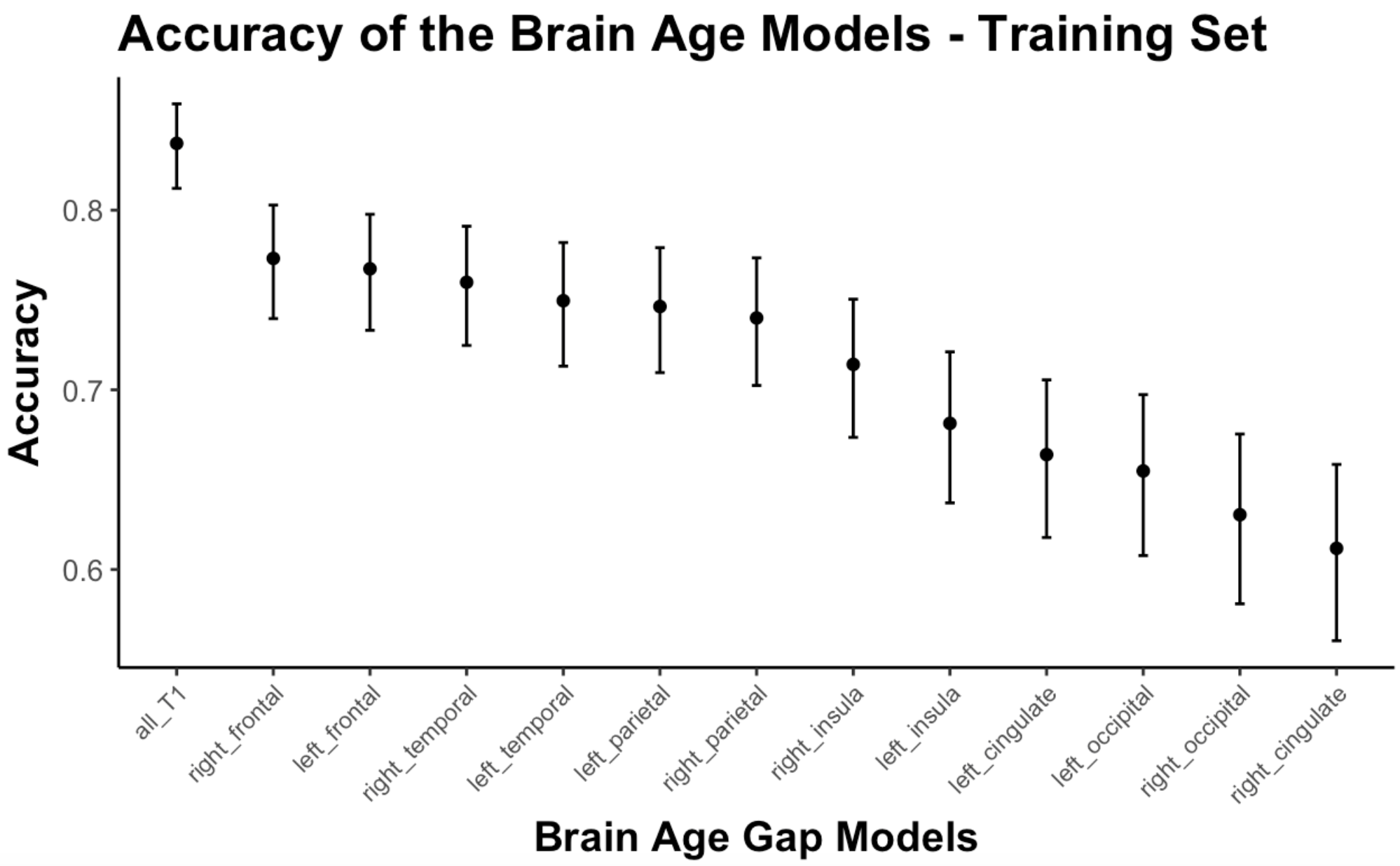

**Suppl. Figure 1.** Pearson correlation with 95% confidence intervals between estimated brain age and chronological age within the training sample.
