## Supplementary material for "Reliable longitudinal brain age prediction in stroke patients: Associations with cognitive function and response to cognitive training": Suppl. Table

Supplementary Tables

| models | *r* | lowerCI | upperCI | MAE |
| --- | --- | --- | --- | --- |
| Global |  |  |  |  |
| all_T1 | 0.550 | 0.358 | 0.697 | 5.506 |
| median | 0.579 | 0.395 | 0.718 | 4.271 |
| Regional |  |  |  |  |
| left_frontal | 0.448 | 0.234 | 0.620 | 5.799 |
| right_frontal | 0.403 | 0.182 | 0.585 | 5.663 |
| left_parietal | 0.413 | 0.194 | 0.593 | 6.287 |
| right_parietal | 0.460 | 0.248 | 0.629 | 6.086 |
| left_occipital | 0.221 | -0.019 | 0.436 | 7.642 |
| right_occipital | 0.206 | -0.034 | 0.423 | 6.953 |
| left_temporal | 0.416 | 0.198 | 0.596 | 7.588 |
| right_temporal | 0.455 | 0.242 | 0.625 | 5.733 |
| left_cingulate | 0.086 | -0.156 | 0.317 | 6.953 |
| right_cingulate | 0.439 | 0.224 | 0.613 | 8.804 |
| left_insula | 0.309 | 0.076 | 0.510 | 7.732 |
| right_insula | 0.366 | 0.140 | 0.556 | 6.277 |

**Suppl. Table 1.** Pearson correlation between estimated brain age and chronological age with their confidence intervals on the test sample at baseline (stroke patients from StrokeMRI sample) for each model, and the MAE calculated from BAGR – After removing outliers.

|  | Baseline (scan 1 and 2) | | | All time points (scan 1 to 3) | | |
| --- | --- | --- | --- | --- | --- | --- |
| models | ICC | lowerCI | upperCI | ICC | lowerCI | upperCI |
| Global |  |  |  |  |  |  |
| BAGR_all_T1 | 0.789 | 0.663 | 0.872 | 0.785 | 0.688 | 0.860 |
| BAGR_median | 0.893 | 0.823 | 0.936 | 0.839 | 0.762 | 0.897 |
| Regional |  |  |  |  |  |  |
| BAGR_left_frontal | 0.789 | 0.664 | 0.872 | 0.811 | 0.723 | 0.878 |
| BAGR_right_frontal | 0.743 | 0.596 | 0.842 | 0.730 | 0.615 | 0.822 |
| BAGR_left_parietal | 0.805 | 0.687 | 0.882 | 0.805 | 0.716 | 0.874 |
| BAGR_right_parietal | 0.894 | 0.825 | 0.937 | 0.856 | 0.786 | 0.909 |
| BAGR_left_occipital | 0.821 | 0.712 | 0.892 | 0.814 | 0.727 | 0.880 |
| BAGR_right_occipital | 0.785 | 0.657 | 0.869 | 0.810 | 0.722 | 0.877 |
| BAGR_left_temporal | 0.796 | 0.674 | 0.876 | 0.804 | 0.713 | 0.873 |
| BAGR_right_temporal | 0.726 | 0.572 | 0.831 | 0.750 | 0.641 | 0.836 |
| BAGR_left_cingulate | 0.663 | 0.484 | 0.789 | 0.695 | 0.571 | 0.797 |
| BAGR_right_cingulate | 0.814 | 0.700 | 0.887 | 0.802 | 0.711 | 0.872 |
| BAGR_left_insula | 0.807 | 0.691 | 0.883 | 0.802 | 0.711 | 0.872 |
| BAGR_right_insula | 0.779 | 0.649 | 0.866 | 0.774 | 0.673 | 0.852 |

**Suppl. Table 2.** Intra-class correlation (ICC) with their confidence interval of the estimated brain age for the two baseline scans (scan one and two), and for the three timepoints (scan one, two and three) – After removing outliers.
